# Targeting Lysyl oxidase-like 2 in Idiopathic Pulmonary Fibrosis

**DOI:** 10.1101/813907

**Authors:** Milena S. Espindola, David M Habiel, Ana Lucia Coelho, Amanda Mikels-Vigdal, Cory M. Hogaboam

## Abstract

The composition of extracellular matrix (ECM) is altered during pathologic scarring in damaged organs including the lung. One major change in the ECM involves the cross-linking of collagen, which promotes fibroblast to myofibroblast differentiation.

**Objective:** We examined the role of lysyl oxidase (LOX)-like 2 in lung fibroblasts cultured from normal or IPF lung samples and in a humanized mouse model of IPF using a monoclonal antibody (Simtuzumab).

**Research Design and Methods:** Primary lung fibroblasts from normal donor lungs and IPF lung explants were examined for expression of LOXL2. Targeting LOXL2 with Simtuzumab on normal and IPF fibroblasts was examined both *in vitro* and *in vivo* for synthetic, functional, and profibrotic properties.

**Results:** LOXL2 was increased at transcript and protein level in IPF compared with normal lung samples. In a dose-dependent manner, Simtuzumab enhanced differentiation of fibroblasts into myofibroblasts. Inhibition of LOXL2 also enhanced fibroblast invasion and accelerated the outgrowth of fibroblasts from dissociated human lung cell preparations. Finally, preventative or delayed delivery of Simtuzumab enhanced lung fibrosis in a humanized mouse model of pulmonary fibrosis.

**Conclusion:** Consistent with its failure in a Phase 2 clinical trial, Simtuzumab exhibited no therapeutic efficacy in translational *in vitro* and *in vivo* assays.

## Introduction

Dynamic remodeling of the extracellular matrix (ECM) is essential for development, wound healing, and normal organ homeostasis but pathological organ fibrosis arises in the context of excessive or uncontrolled ECM remodeling ^1^. Lysyl oxidase (LOX) and LOX-like (LOXL) proteins (i.e. LOX family-LOX, LOXL1, LOXL2, LOXL3 and LOXL4) play crucial roles in ECM remodeling including crosslinking of collagens and elastin due and LOXL2 protein was identified as a secreted protein and localized in the extracellular matrix in active fibrotic diseases and in the early stromal reaction of breast cancer ^2^. Subsequent studies showed that LOXL2 expression was increased in the context of several fibrotic conditions including those in the liver ^3, 4, 5^, heart ^6, 7^, cancer ^8^, eye ^6, 9^, and lung ^10^.

In the lung, ECM is a key component of airways and lung parenchyma providing physical support and stability to these areas in the lung. Alterations in the ECM are evident in several chronic lung diseases including asthma, chronic obstructive pulmonary disease (COPD), and idiopathic pulmonary fibrosis (IPF), in which the folding or rigidity of ECM proteins changes the functions of key cell types including fibroblasts^11, 12^. IPF is a progressive and lethal disease characterized by excessive deposition of ECM in the lung parenchyma ^13^. Constitutive expression of all LOX/L family members was detected in normal cells such as lung epithelial cells and fibroblasts, and the latter cell type exhibited the induction of LOXL2 following exposure to pro-fibrotic stimuli ^10^. In IPF lung tissues, LOX and LOXL2 was detected in bronchial and alveolar epithelium as well as fibroblastic foci using immunohistochemical approaches ^10^. Elevated sLOXL2 levels in serum were also associated with increased risk for IPF disease progression in two separate cohorts of IPF patients ^14^. Allosteric inhibition of LOXL2 with Simtuzumab inhibits the enzymatic function of LOXL2 in a non-competitive manner thereby allowing inhibition of LOXL2 regardless of substrate concentration ^15^. This unique mAb showed impressive efficacy in experimental models of pancreatic cancer, liver fibrosis, lung fibrosis ^16^, cardiac fibrosis ^6^, and choroidal neovascularization ^17^ but this targeting approach failed in Phase 2 clinical trials directed at the first three conditions. Details regarding the clinical trial in IPF were reported by Raghu et al. and the conclusion drawn from that study was that Simtuzumab did not improve progression-free survival in a well-defined population of patients with IPF ^18^.

Since the failure of Simtuzumab in this IPF trial was unexpected given the convincing preclinical data generated in mouse, we undertook both *in vitro* and *in vivo* studies to address the efficacy of Simutuzumab in translational studies. Specifically, we examined the effect of this mAb in primary human fibroblasts from both normal lung donor samples and explanted IPF lung samples and in a humanized SCID model of IPF^19^. Overall, our data suggested that targeting the cross-linking of ECM with Simtuzamab promoted IPF fibroblast to myofibroblast transition and increased IPF fibroblast invasion. In the *in vivo* model, mAb delivery either in a preventative or therapeutic manner did not alter the fibrotic response evoked by the intravenous injection of IPF lung cells into SCID mice. Together, these data highlight the importance of both *in vitro* and *in vivo* translational studies with human cells to provide key assurances of target validation in IPF.

## Methods

### Patients and Study approval

Institutional Review Boards both at Cedars-Sinai Medical Center (Pro34067) and the University of Michigan (Hum00004350) approved all experiments with primary human tissue and mouse studies described herein. All patients were consented prior to inclusion in the studies.

### Gene expression array data mining and Ingenuity IPA analysis

Publicly available gene expression datasets (GSE24206) were mined from NCBI’s geo datasets database. Groups were defined as follows – IPF lung biopsies vs normal lungs and IPF lung explants vs normal lungs. Gene expression values were extracted using NCBI’s Geo2R gene expression analysis tool and the expression data were uploaded onto ingenuity IPA. Ingenuity IPA was set to only consider changes in gene expression of 1.5-fold or greater and p ≤ 0.05. The resulting analysis was then overlaid onto a custom LOXL2 interaction network, generated using Ingenuity’s knowledge database for all proteins that have been reported to directly interact with LOLX2.

### Immunohistochemistry

Slides containing 4 µm sections of IPF human lung biopsies or explants were deparaffinized and hydrated by incubating them twice in xylene for five min each, followed by 2 changes of 100% ethanol for 3 min each, 70% ethanol for 2 min, 50% ethanol for 2 min, and distilled water for 5 min. Antigen retrieval was performed by incubating the slides in 10 mM citric acid solution (pH 6.0) in an 80 °C oven overnight. The slides were then washed in PBS and permeabilized in 10% methanol containing 0.4% H_2_O_2_ for 30 min. After permeabilization, slides were blocked and stained using rabbit anti-LOXL2 (Gil25-70, Gilead Sciences) antibody overnight at 4° C and a rabbit cell & tissue staining kit (R&D Systems) as recommended by the manufacturer. Immuno-stained lung tissues were images were obtained at 20*x* and 40x magnification.

### Isolation of mixed cells from IPF explants

Normal and IPF lung explants were acquired from consented donors. Fresh explant cells were isolated as previously described ^19^.

### Isolation and Culture of Primary Pulmonary Fibroblast Lines

IPF surgical lung biopsy samples were obtained as previously described ^20^. IPF lung fibroblasts were generated by mechanically dissociating IPF lung biopsies or explants as previously described ^21^.

### Protein analysis

Lung fibroblasts (0.5 × 10^4^/well) were plated into a 96 well plate and incubated overnight. After incubation, cells were treated with anti-LOXL2 (1, 10, and 50 μg/ml, GS-6624, Gilead Sciences), BIBF1120 (300 nM) or IgG in the absence or presence of CpG (10 nM), IL-13 (10 ng/ml) or TGF-β (20 ng/ml) for 24 for αSMA expression or 7 days for Collagen 1 assessment. After stimulation, conditioned supernatants were collected and the cells were washed and fixed with 4% paraformaldehyde solution in PBS for 10 min at room temperature.

#### In cell αSMA

After fixation, cells were washed and permeabilized with 0.5% Triton X-100 in DPBS for 10 min, washed and endogenous peroxidase activity was blocked by adding a solution of 0.3% H_2_O_2_ in DPBS to the cells and incubating the cells with the solution for 20 min at room temperature. Cells were then washed and wells were blocked with a 1% BSA solution in DPBS for 30 min at room temperature. After blocking, cells were incubated with 500 ng/ml of anti-αSMA (Clone 1A4, Abcam) antibody overnight at 4 °C on a rocker. After incubation, cells were washed and incubated with an HRP conjugated rabbit anti-mouse antibody (1:5000 diluted, R&D systems) for 30 min at room temperature. Cells where then washed and the reaction was developed using 100 µl of TMB developing reagent (Fitzgerald Industries International) until sufficient color was observed. The reaction was then stopped by adding 50 µl of 2N sulfuric acid and data were acquired by reading the absorbance at 405 nm using a Synergy H1 microplate reader (BioTek Instruments Inc.). To normalize to cell number, the plate was then washed, and cells were incubated with 500 ng/ml of anti-β-tubulin antibodies (Abcam) and incubated overnight at 4 °C. Cells were then washed, incubated with 200 ng/ml of AP-conjugated goat anti-rabbit IgG antibody (SeraCare Life Sciences) for 1 h at room temperature, then washed and developed using a Bluephos AP developing reagent (SeraCare Life Sciences). Data were acquired by reading the absorbance at 595 nm using a Synergy H1 microplate reader (BioTek Instruments Inc.). Ratios were generated from αSMA/β-tubulin absorbance data and the results were normalized to vehicle. The fold change after 24 relative to IgG was then determined and depicted in the figures.

#### Soluble Collagen 1 ELISA

Lung fibroblast conditioned supernatants were diluted 1:5 in DPBS. Purified Collagen 1 (STEMCELL Technologies Inc), diluted in 1:5 DBPS diluted complete medium, was utilized to generate an 8-point collagen 1 standard ranging from 200-0 ng/ml. Fifty microliters of the supernatants and standards were coated onto multisorb 96 well plates (Thermo Fisher Scientific) overnight at 4 °C. After coating, the plates were washed and blocked with 200 µl of 3% BSA solution in DPBS (Lonza) for 1 h at room temperature on an orbital shaker. After blocking, plates were washed and 50 µl of 200 ng/ml biotin-conjugated anti collagen-1 antibody (Ab2482,1Abcam) was added to each well and the plates were incubated for 2 h at room temperature on an orbital shaker. After incubation, plates were washed and 50 µl of HRP-conjugated streptavidin (R&D systems, diluted as recommended by the manufacturer) was added to each well and the plates were incubated for 30 min at room temperature on an orbital shaker. Plates were then washed and the wells were developed by adding 100 µl of TMB peroxidase substrate solution (Fitzgerald Industries International) until sufficient color is observed, after which the reaction was stopped by adding 50 µl of 2N Sulfuric acid.

### *In vitro* fibroblast invasion

For fibroblast scratch wound invasion, 96 well ImageLock™ plates (Essen BioScience) were coated with 50μg/mL of Basement membrane extract (BME) (Trevigen) for a minimum of one hour at room temperature Normal and IPF fibroblasts were added in a concentration of 3.5 × 10^4^/well overnight. Cells were then scratched using the WoundMaker™ (Essen BioScience), washed with PBS and then incubated with the respective treatments added with BME (4mg/mL) in complete medium for a least 96 hours at 37 °C, 10% CO_2_. Wound closure was measured as percentage of closure, quantitated via IncyCyte ZOOM software (Essen Biosciences) as recommended by the manufacturer.

### Humanized mouse model of pulmonary fibrosis

Six-to 8-week-old, female, pathogen-free NOD Cg-PrkdcSCID IL2rgTm1wil Szi (NSG) were purchased from Jackson Laboratories and housed in Cedars-Sinai Medical Center’s high isolation mouse room. NSG mice were allowed a minimum of 1 week to recover from transport in the facility and then each mouse received nothing (i.e., naïve or non-humanized) or approximately 1 × 10^6^ IPF cells isolated from explanted lung samples. IPF cells were previously bio-banked in liquid N_2_ and prior to intravenous injection into NSG mice, these cells were thawed, washed in serum-free medium, and suspended in a volume of 0.5 ml. Treatments with either IgG (AB005123, GS645864, Gilead Sciences) or anti-LOXL2 (AB0023, GS607601, Gilead Sciences) mAb from day 0-35 (i.e. preventative regimen) or from day 35-63 (i.e. therapeutic regimen) after intravenous injection of IPF lung cells. Sixty-three days after IPF cell injection and IgG or anti-LOXL2 mAb treatment, the superior and middle lobes were collected for biochemical hydroxyproline quantification, the post-caval lobe for qPCR analysis, and the left lung for histological analysis from each NSG mouse.

### Hydroxyproline assay

Total lung hydroxyproline was analyzed as previously described ^21^.

### Quantitative PCR analysis

Cells or lung lobes were lysed in Trizol™ reagent. RNA was extracted as recommended by the manufacturer and 1 µg of RNA was reverse transcribed into cDNA using superscript II reverse transcriptase (Life technology) as previously described ^20^. Complementary DNA (cDNA) was subsequently loaded into a TaqMan plate (Thermo-Fisher Scientific) and gene expression analysis were performed using predesigned primers and probes for human-*LOXL2, ACTA2, COL1a1, COL3a1, FN1, RNA18S5* and mouse-*col1a1, col3a1, fn1, tgfb, pdgfra and pdgfrb* (Thermo-Fisher Scientific). All TaqMan analysis was performed using an Applied Bio system’s Viia 7 instrument (Thermo-Fisher Scientific). The results were then exported, normalized to *RNA18S5* (Thermo-Fisher Scientific) and fold change values were calculated using DataAssist software (Thermo-Fisher Scientific).

### Histological analysis

Mouse lung tissue was fixed in 10% Neutral buffered formalin (NBF) solution for 24 h and subsequently transferred into tissue cassettes and kept in a 70% ethanol solution for approximately 24 h. Lungs were then processed using routine histology techniques and stained using Masson’s trichrome. Images were acquired using the Leica Slide Scanner SCN400 and SCN400 client software (Leica Microsystems Inc., SL801). The digital images were documented by exporting to the Digital Image Hub (DIH-SlidePath).

## Results

### LOXL2 transcript levels are elevated in both IPF lung biopsies and lung explants compared with normal lung donor samples

We first examined the transcriptional landscape of LOXL2 using Ingenuity Pathway Analysis or IPA, which is a web-based software application for the analysis of data derived from various omics platforms but more heavily weighted toward RNA. Analysis of publicly available gene arrays revealed that LOXL2 was strongly upregulated in both diagnostic biopsies and in explanted lung tissues, and several downstream factors were also upregulated in a statistically significant manner compared with normal lung samples (**Fig. 1**). FN1, VIM, MARCKSL1, SMARC4, MTA1 were the downstream factors that were consistently increased in early (i.e. diagnostic biopsy) (**Fig. 1A**) and late (i.e. lung explant) (**Fig. 1B**) IPF compared with normal lung samples. Thus, our data were consistent with previous reports in which it was observed that LOXL2 transcript levels were increased in IPF ^10, 22^.

**Figure 1.**
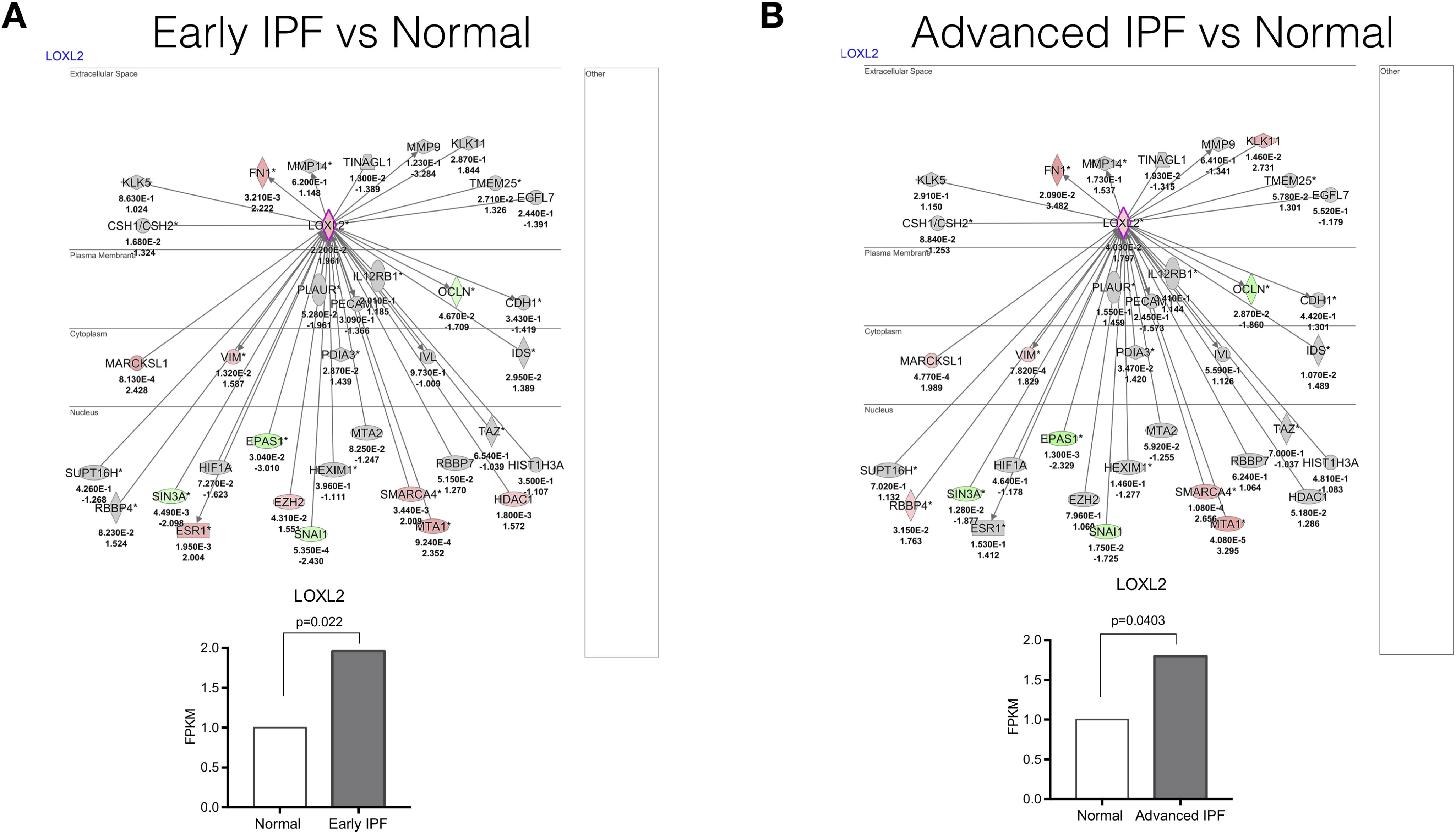
LOXL2 exhibits increased expression in active disease. An LOXL2 interaction network was generated using Ingenuity’s knowledge base and overlaid on relative expression datasets from lung tissue microarray datasets. (A and B) P values (top) and fold change in expression (IPF vs. normal; bottom) from publicly available datasets (GSE24206) comparing IPF surgical lung biopsies (A) or IPF explants (B) to normal lung tissues. Up- and downregulated transcripts are depicted in red and green, respectively.

### LOXL2 protein is present in both IPF lung biopsies and lung explants

To confirm the transcriptional analyzes, we next used immunohistochemical approaches to localized LOXL2 in both diagnostic biopsies and explanted lung. As shown in **Figure 2**, LOXL2 was strongly upregulated in cells in many cell types in the diagnostic biopsy (**Fig. 2A**) but the expression of LOXL2 was almost exclusively extracellular in explanted lung samples (**Fig 2B**). The latter finding was consistent with previous findings that this enzyme is secreted and predominately localizes to the ECM in the IPF lung ^22^.

**Figure 2.**
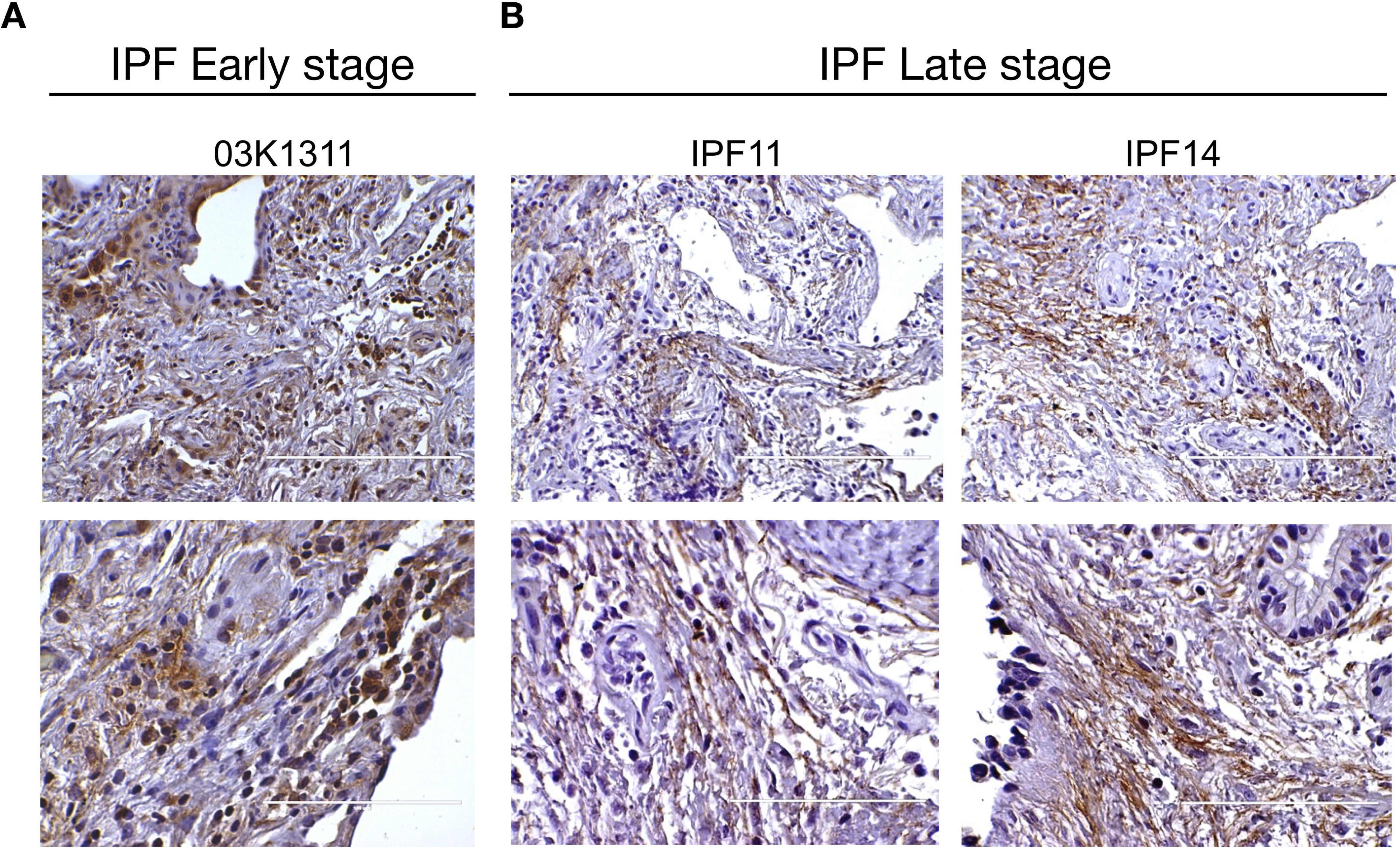
LOXL2 expression on IPF tissue. Representative LOXL2 expression in immunohistochemical sections from (**A**) early disease stage (biopsies) and (**B**) late disease stage (center and right panels). Top and bottom panel images were obtained at 20x or 40x magnification, respectively.

### Targeting LOXL2 with Simtuzumab enhances fibroblast to myofibroblast differentiation and fibroblast invasion *in vitro*

Next, we utilized the cells obtained from normal or IPF lungs in a series of *in vitro* studies with cultured primary fibroblasts from normal donor lung samples and IPF lung explants. LOXL2 transcript expression was present in both normal and IPF lung fibroblasts although levels of LOXL2 were modestly increased in IPF fibroblasts relative to normal fibroblasts (**Fig. 3A-B**). Mixed lung explanted cells expressed significantly less LOXL2 compared to fibroblasts, although IPF cells showed modest increased expression compared to Normal cells. (**Fig. 3C-D**). The dose-dependent effect of Simtuzumab on the expression of alpha smooth muscle actin protein in both normal and IPF fibroblasts was examined using an in-cell ELISA we developed in the laboratory. As indicated in **Figure 4A**, various stimuli were used to alter alpha SMA including hypomethylated DNA (or CpG), IL-13, or TGF-β (see Figure legend for concentrations of each stimuli). In cultures of normal human lung fibroblasts, the doses of stimuli used were not effective in the induction of alpha SMA (i.e. myofibroblast differentiation) unless 50 micrograms/ml of Simtuzumab was present or 300 nM of BIBF-1120 (also known as nintedanib). Note that the dose of 50 micrograms/ml was shown previously to effectively prevent collagen crosslinking ^16^. In IPF fibroblasts, the stimuli were more effective at driving myofibroblast differentiation, and both anti-LOXL2 mAb treatment and BIBF-1120 enhanced myofibroblast differentiation. Soluble collagen 1 was quantified on supernatants of fibroblasts treated with 50 micrograms/ml of Simtuzumab or BIBF-1120 under the stimuli previously described. Simtuzumab failed in altering or reducing collagen synthesis by these cells, as opposed to BIBF-1120 that significantly reduced collagen 1 production by fibroblasts treated with TGF-β (**Fig. 4B**). In a scratch wound assay, BIBF-1120 effectively prevented fibroblast migration whereas the presence of Simtuzumab at the 50-microgram dose increased the speed of fibroblast migration compared with the IgG control condition (**Fig. 4C**). All together, these in vitro data suggest that targeting LOXL2 in both normal and IPF primary lung fibroblasts did not alter myofibroblast differentiation or the invasive properties of these cells in a well-established scratch wound invasion assay.

**Figure 3.**
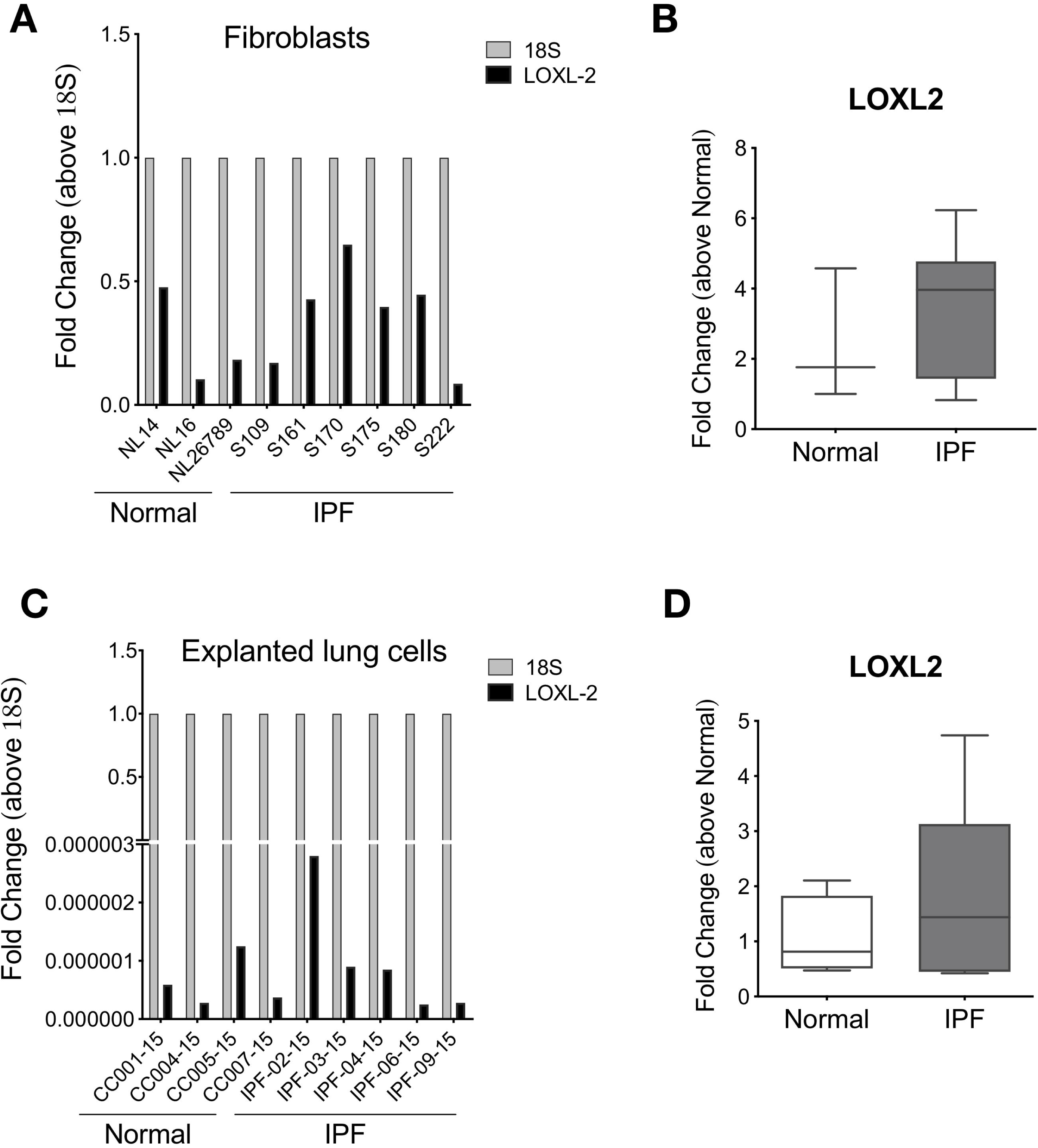
Fibroblasts and lung explanted cells express increased LOXL2. Fibroblast cultures derived from normal and IPF lung samples or total lung explanted cells were submitted to quantitative PCR for LOXL2. (A) Relative expression of LOXL2 in Normal and IPF Fibroblast cell lines and (B) relative fold change compared to Normal fibroblasts. (C) Relative expression of LOXL2 in Normal and IPF total explanted cells from IPF lungs and (B) relative fold change compared to Normal explanted lung cells.

**Figure 4.**
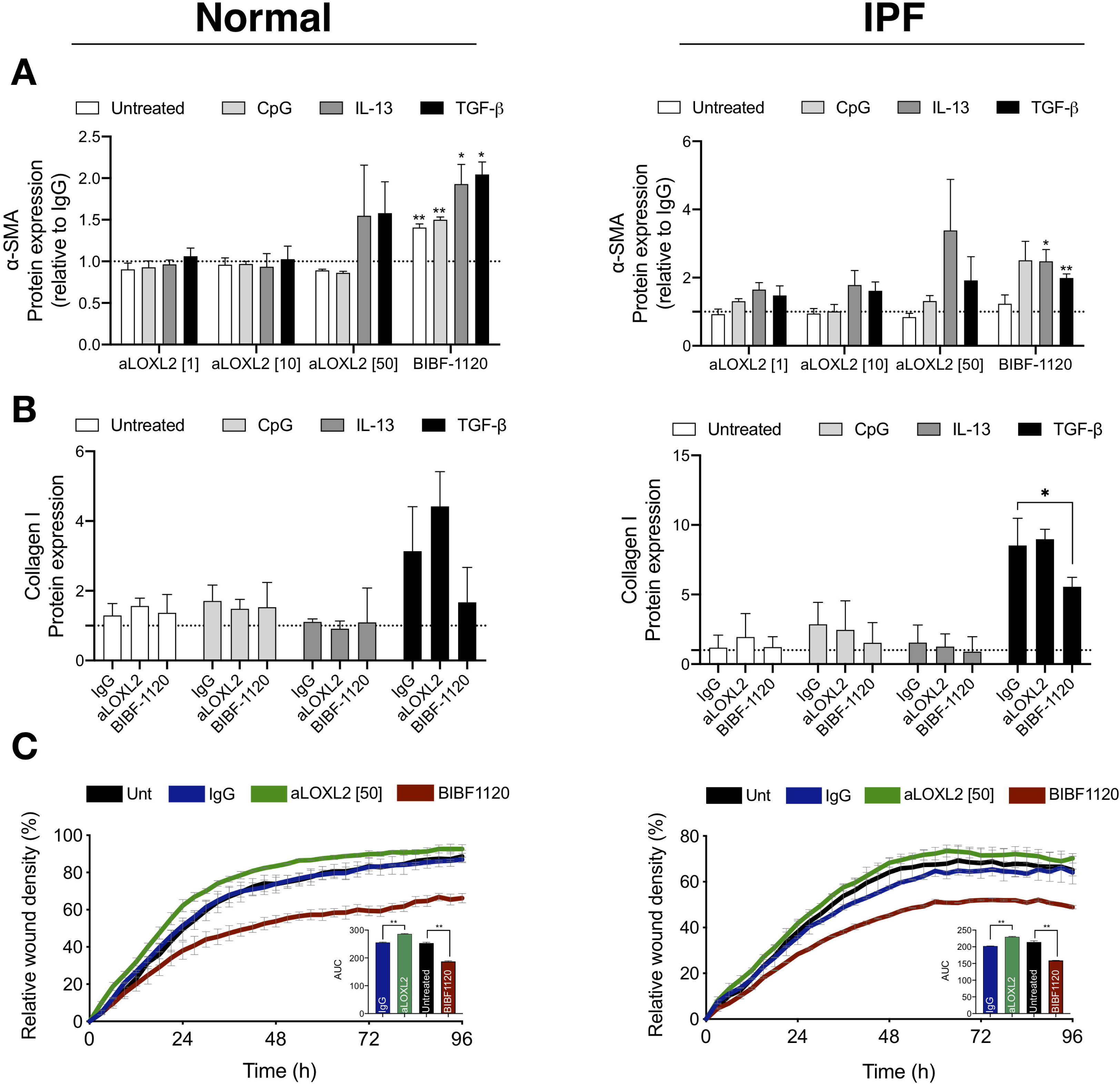
Effects of Simtuzumab on normal and IPF myofibroblast differentiation and invasion. (**A-B**) Fibroblast cultures derived from normal (left panels) and IPF (right panels) lung samples were treated with anti-LOXL2 (1, 10, and 50 μg/ml), BIBF1120 (300 nM) and their respective controls in the absence or presence of CpG (10 nM), IL-13 (10 ng/ml) or TGF-β (20 ng/ml). (**A**) α-SMA was quantified via *in situ* in-cell ELISA 24h after stimulation and results are expressed relative to IgG. (**B**) Collagen 1 was measured via direct ELISA 7 days after treatment with anti-LOXL2 (50 μg/ml), BIBF1120 (300 nM) or IgG. Results are expressed relative to Untreated IgG samples. (**C**) Functional analysis of the role of LOXL2 in fibroblast invasion. Normal (left panel) and IPF (right panel) fibroblasts were left untreated or treated with anti-LOXL2 (50 μg/ml), BIBF1120 (300 nM) or IgG. Invasive properties were assessed by scratch wound invasion for a minimum of 96h. The area under the curve for each treatment is depicted on the bar graph of representative cell lines (n=2-3). *P ≤ 0.05; **P ≤ 0.01 compared to IgG or Untreated samples.

### Outgrowth of primary fibroblasts from lung cells dissociated from either normal donor lungs or IPF lung explants

Our next *in vitro* study addressed how collagen crosslinking affected the outgrowth of fibroblasts from mixed human lung cells added to tissue culture flasks. In this study, mixed cells from either normal donor lungs or IPF lung explants were seeded at defined cell concentrations and the cultures were monitored for the presence of fibroblasts at days 7, 14, 21, and 26. At all timepoints examined, fibroblasts were more abundant in cell cultures supplemented with Simtuzumab compared with cell cultures supplemented with the appropriate IgG control (**Fig. 5A-B**). This accelerated outgrowth was confirmed using qPCR analysis for various fibrosis-related genes including smooth muscle actin (ACTA2), collagen 1a1 (COL1a1), collagen 3a1 (COL3a1), and fibronectin (FN1) at day 26 after the start of this experiment for both groups. The levels of all transcripts, except COL3a1 on IPF cells, were increased in both normal and IPF cultures treated with the anti-LOXL2 antibody over the course of the culture time (**Fig. 5C**). Thus, the presence of Simtuzumab appeared to accelerate the outgrowth of both normal and IPF fibroblasts from dissociated lung samples.

**Figure 5.**
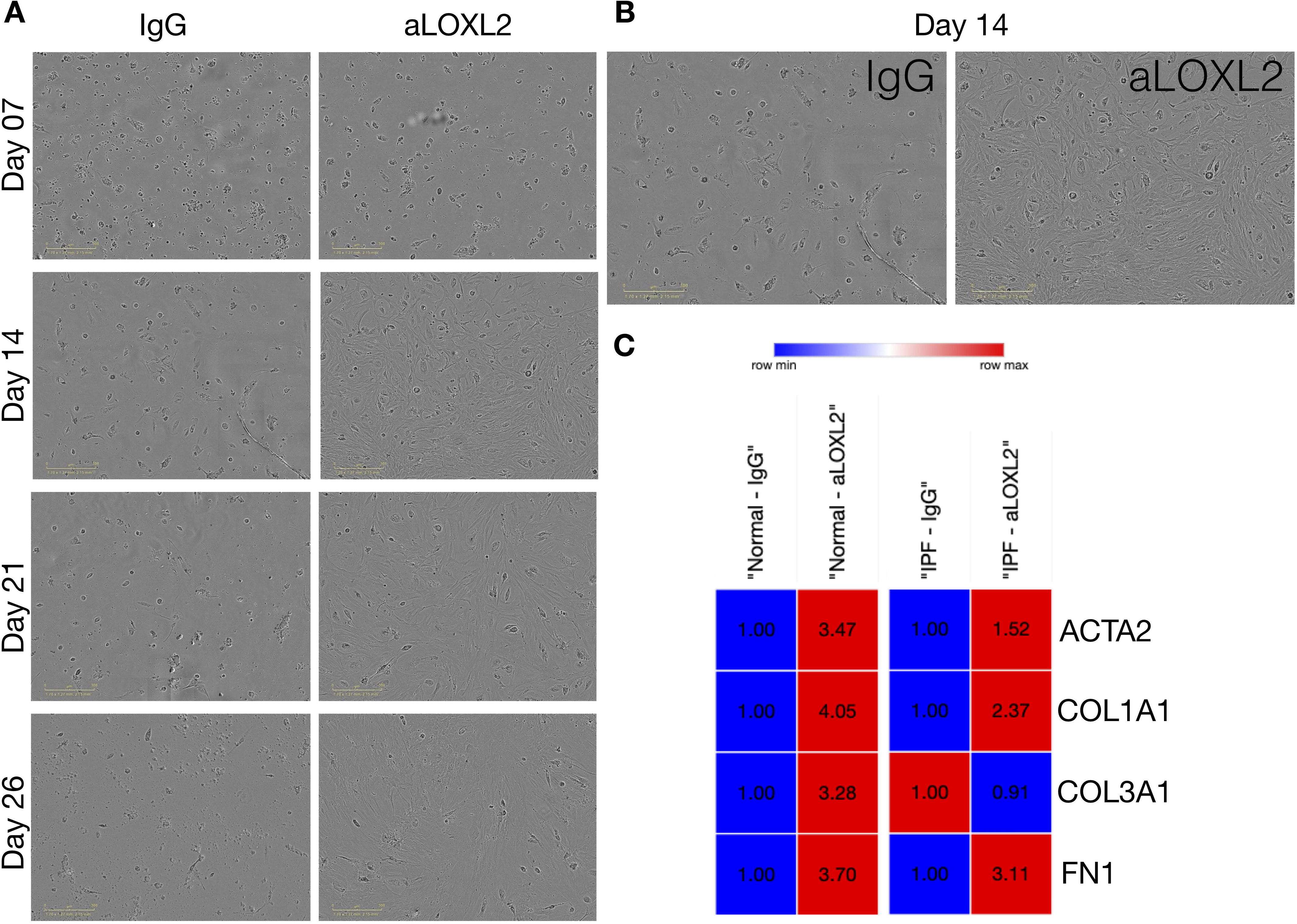
Simtuzumab accelerates cell differentiation *in vitro*. Explanted lung cells from normal and IPF lung samples were treated with anti-LOXL2 (50 μg/ml), or IgG for 26 days. Media and treatments were replaced 2x/week until day 26. (**A**) Representative images from cell cultures were collected at day 07, 14, 21 and 26. (**B**) Detailed images from Day 14 showing mesenchymal cells differentiation on LOXL2 treated cells. (**C**) Heatmaps from Normal and IPF cell cultures for ACTA2, COL1A1, COL3A1 and FN1. Transcripts were assessed by quantitative PCR (n=2).

### Preventative or therapeutic targeting of LOXL2 fails to alter lung fibrosis in a humanized SCID mouse model of IPF

Our last study addressed the role of LOXL2 *in vivo* using a humanized SCID model of IPF in which approximately half a million freshly isolated mixed cell populations from explanted IPF lungs are injected intravenously into SCID mice to initiate the lung fibrotic process ^19^. In this model, fibrosis appears in the lung approximately 30-35 days after human cell injection and the fibrosis persist beyond the day 63 timepoint we typically end our studies ^19^. In our present study, we employed a preventative treatment modality that involved the injection of 15 mg/kg of IgG or Simtuzumab twice a week up to day 35. In a separate therapeutic study, injections of either IgG or Simtuzumab at 15 mg/kg twice a week began at day 35 and continued up to the termination of the study at day 63. In the preventative treatment groups, anti-LOXL2 mAb treatment significantly increased lung hydroxyproline levels in humanized SCID mice that received human cells versus mice that received human cells alone (**Fig. 6A, left panel**). However, there was no difference in the Simtuzumab-treated group compared with the IgG-treated group. In the therapeutic treatment groups, humanized SCID mice treated with Simtuzumab had a statistically significant increase in total lung hydroxyproline compared with naïve SCID mice (**Fig. 6A, right panel**). Again, no differences were observed between the Simtuzumab- and IgG-treated groups of humanized SCID mice. Transcript analysis of whole lung samples from the day 63 groups showed that there were no changes in fibrosis-related transcripts in humanized SCID mice treated with either IgG or with anti-LOXL2 mAb (**Fig.6B**) and the histological picture at this time point was similar between the two groups of humanized SCID mice regardless of when the treatment was administered (**Fig. 6C-D**). Taken together, targeting LOXL2 did not modulate the fibrotic response in this translational model of IPF regardless of the treatment modality.

**Figure 6.**
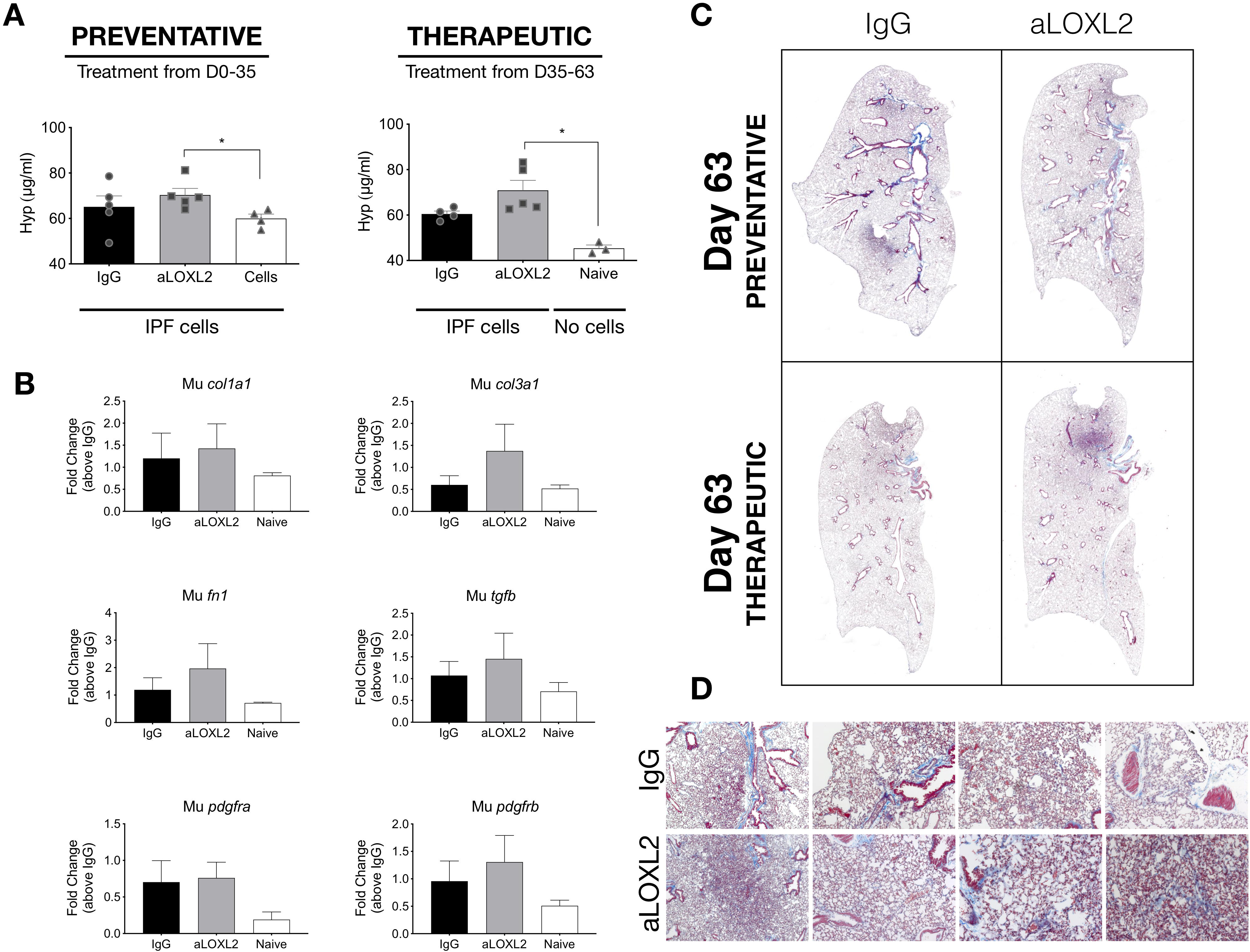
Effects of Simtuzumab in experimental pulmonary fibrosis. Cellular suspensions of IPF lung explants were injected intravenously into NSG mice and allowed to engraft for 35 days. Humanized NSG mice were treated with IgG or anti-LOXL2 (15mg/kg) in a Preventative or Therapeutic manner, twice a week from days 0 to 35 or from days 35 to 63, respectively. Mice were sacrificed, and their lungs were analyzed for remodeling. (**A**) Hydroxyproline content in the superior and middle lobes of NSG mouse lungs; (**B**) Transcript expression of murine col1a1, col3a1, fn1, tgfb, pdgfra and pdgfrb (**C**) Representative images of Masson’s trichrome staining in whole lungs and (**D**) affected areas at 20x magnification of NSG mouse lungs at day 63 after IPF cell administration. (n=3-5). *P ≤ 0.05 where indicated.

## Discussion

Cross linking of collagen fibers in the ECM is a key hallmark of progressive and unrelenting fibrotic conditions including IPF ^23^. There are several enzymes that appear to be involved at various stages of collagen assembly in the lung and other organs, consequently it has proven to be very challenging to identify effective targeting strategies that modulate ECM remodeling appropriately. LOXL2 emerged as one of the most markedly and consistently altered catalysts for the formation of cross-links in fibrillar elastin and collagens thereby generating considerable attention in an ECM-associated pathology such as IPF ^23^. Studies in a bleomycin-induced pulmonary fibrosis model strongly suggested that LOXL2 was an attractive target in fibrosis ^16^ and lead to a clinical trial designed to investigate the efficacy and safety of Simtuzumab, a monoclonal antibody against LOXL2, in patients with IPF ^18^. The failure of Simtuzumab to modulate the course of disease in IPF patients motivated us to examine more closely how LOXL2-mediated collagen crosslinking impacted the synthetic and functional properties of primary normal and IPF lung fibroblasts both *in vitro* as well as in a humanized SCID model of IPF. Overall, we observed that targeting LOXL2 enhanced the synthetic and invasive properties of both primary normal and IPF fibroblasts and accelerated the outgrowth of these cells from dissociated cells from normal or IPF lung samples. The targeting of LOXL2 *in vivo* further indicated that this enzyme was not required for the development of pulmonary fibrosis in this model. Together, our findings are a cautionary tale regarding the importance of translational studies in IPF target validation.

Collagen cross-linking mediated by the LO family of enzymes, most notably LOXL2, has been examined in various in vitro studies and the role of these enzymes in this process appears to be firmly established. Specifically, previous studies had indicated that limiting the process of fibrillar collagen cross-linking was likely to correct the pathological microenvironment present in IPF ^22^. In the present study, we confirmed that LOXL2 was increased in IPF compared with normal lung samples and cells, but the reduction in collagen cross-linking with Simtuzumab clearly enhanced the synthetic and functional activity of both normal and IPF fibroblasts in the assays we explored. The reason for the apparent discrepancy between previous studies and our own are not clear at the moment but we are confident that the fibroblasts we studied were responsive to anti-fibrotic therapy as evidenced by the effects of BIBF-1120 on these cells. The outgrowth of fibroblasts from mixed cell cultures was most surprising since the expectation in this system is that LOXL2 would have little effect on this response. Adding fibroblasts directly to tissue culture plastic can alter the behavior of these cells in ways that are beyond that observed in pathologic tissue conditions, but we did include studies in which primary fibroblasts were mixed into Matrigel and were not in contact with tissue culture plastic during the course of the scratch wound assay. Thus, the translational in vitro studies we undertook did not indicate that targeting LOXL2 was anti-fibrotic but instead indicated that cross-linked collagen might have a modulating effect on the activation of these cells.

Previous studies in humanized SCID mouse models of lung cancer induced with the introduction of Lewis lung carcinoma (LLC) cells suggested that Simtuzumab was beneficial for the treatment of angiogenic tumors ^24^. In our study using a humanized SCID mouse model to examine pulmonary fibrosis, we did not observe either a preventative or therapeutic effect of Simtuzumab. The introduction of primary lung cells from IPF explants initiates and sustains a fibrotic response in the lungs of humanized SCID mice and we have observed that this model is responsive to anti-fibrotic therapies ^19^, although Nintedanib showed no therapeutic effect in the humanized SCID IPF fibroblast model ^21^. The failure of Simtuzumab in this model was unlike that observed in the bleomycin-induced pulmonary fibrosis model^16^, and highlights the need for multiple model testing before moving a target into the clinic.

Presumably, several explanations account for the failure of Simtuzumab in IPF, and many of these are emerging as new research proceeds in the area of ECM remodeling. For example, exploration of epigenetic mechanisms that regulate fibroblast activity in cancer point to the role of miR-29a and its direct effect on both LOXL2 and serpin peptidase inhibitor clade H, member 1 (SERPINH1) ^25^. These studies suggested that both enzymes worked in tandem to alter collagen synthesis and modification. In addition, other studies point to other non-enzymatic functions of LOXL2 alter cell behavior so that that agents that inhibit the enzymatic activity of LOXL2 might not completely negate the effects of LOXL2 ^26^. In our studies, we did not explore these putative reasons for the failure of Simtuzumab to modulate the pro-fibrotic properties of primary human fibroblasts both *in vitro* and *in vivo*. Future studies are certainly warranted to address the compensatory mechanisms that account for the lack of efficacy of Simtuzumab in IPF.

LOXL2 catalyzes collagen and elastin to remodel the ECM and there is nothing in this current study to refute the importance of this process in IPF. With the clinical failure of Simtuzumab it would appear that there are several challenges around the targeting of ECM modulating enzymes such as LOXL2 in IPF. Emerging evidence suggest that dual LO/LOX targeting approaches are more effective that LOXL2-specific targeting in fibrotic conditions around tumors ^27^. In addition, other studies revealed the important synergistic roles of cellular contractility (via relaxin) and tissue stiffness (via LOXL2) in the maintenance of fibrosis suggesting that these two inter-related mechanisms should be targeted concomitantly ^28^. Finally, LOX is emerging as a potentially useful companion biomarker for evaluation of early on-target effects of other anti-fibrotic agents ^29^. Our translational studies using both normal and IPF cells highlight that targeting the cross-linking of collagen alters properties of primary lung fibroblasts that at least *in vitro* appeared to enhance the functional and synthetic properties of these cells. These findings coupled with the lack of efficacy of Simtuzumab in a humanized SCID mouse model of pulmonary fibrosis highlight the need to further exploration of an IPF target beyond that of a bleomycin-induced pulmonary fibrosis model before proceeding to the clinic. Regardless of the issues associated with targeting LOXL2 in pulmonary fibrosis, caution is certainly warranted as Biotech and Pharmaceutical Companies proceed with other targeting approaches toward LO and LOX and simply rely on preclinical data from bleomycin-induced pulmonary fibrosis models ^30^.

## Acknowledgements

This study was supported by funds and reagents from Gilead Sciences and Cedars-Sinai Medical Center. The authors would like to also acknowledge Kathy McClinchey and McClinchey Histology Lab Inc. for the histological service.

## Notes

The authors have declared that no conflict of interest exists.

